# The Hippocampus Rapidly Integrates Sequence Representations During Novel Multistep Predictions

**DOI:** 10.1101/2025.09.15.676324

**Authors:** Hannah Tarder-Stoll, Christopher Baldassano, Mariam Aly

## Abstract

Memories for temporally extended sequences can be used adaptively to predict future events on multiple timescales, a function that relies on the hippocampus. For such predictions to be useful, they should be updated when environments change. We investigated how and when new learning shapes hippocampal representations of temporally extended sequences, and how this updating relates to flexible predictions about future events. Human participants learned sequences of environments in immersive virtual reality. They then learned novel environment transitions connecting previously separate sequences. During subsequent fMRI, participants predicted multiple steps into the future in both the newly connected sequence and control sequences that remained separate. The hippocampus integrated representations of the connected sequence, such that activity patterns became more similar across trials for the connected sequence vs. the unconnected sequences. These integrated sequence representations in the hippocampus emerged soon after learning, incorporated representations of the initial sequences as well as new activity patterns not previously present in either sequence, and predicted participants’ ability to update their predictions in behavior. Together, these results advance our understanding of how structured knowledge dynamically emerges in service of adaptive behavior.

## Introduction

Memories guide adaptive behavior by supporting predictions. Through repeated experience, the brain – particularly the hippocampus and surrounding medial temporal lobe cortex – creates predictive models that capture the temporal structure of the environment and support future-oriented behaviors at multiple timescales [1–3]. A common challenge, however, is that the environment’s structure does not always remain stable. New information should cause us to update internal models of the environment to support flexible predictions about novel scenarios. For example, learning routes through a city allows us to plan trajectories multiple steps ahead. Upon learning that two seemingly separate routes are linked via a side street, we can plan trajectories along the connected route, even without direct experience [4–8]. Here, we investigated *when* and *how* new learning that connects previously separate, temporally extended sequences shapes hippocampal representations to support flexible prediction of novel trajectories.

The hippocampus is well situated to support flexible prediction because it represents temporal structure and integrates information learned across discrete episodes. For example, hippocampal sequence representations allow individuals to generate multistep predictions about future states [1,3,9,10]. Learning sequential structure requires forming connections among events, and indeed, the hippocampus supports integration across related experiences by linking episodes that share overlapping elements, even if these elements were not encountered close to one another in time [11–16]. In this way, flexible hippocampal representations build structured knowledge that generalizes beyond specific episodes, enabling inferences about novel relationships [16–18]. Yet, prior work has largely examined integration of simple, pairwise associations [11,12,19–27], and real-world environments often involve temporally extended event sequences. In these contexts, new information that links two previously unrelated sequences can have broader consequences, potentially reshaping neural representations of the entire sequential structure even if the change occurs at a relatively local point. Open questions therefore remain about when integrative representations of sequences emerge, what information they contain, and how they influence prediction updating. Here, we investigated both the *time course* and *nature* of hippocampal representations that allow integration to update internal models of temporally extended experiences to guide future-oriented behavior.

We first focus on the time course of updating. In behavior, even brief learning episodes trigger rapid reconfiguration of structured knowledge [28,29]. However, it remains unclear whether new learning coincides with rapid updating of hippocampal representations, or whether these representations are reshaped gradually over time. The hippocampus is well-positioned to support rapid flexibility: it quickly encodes new associations [30,31] which may allow rapid incorporation of new information during planning [32]. However, updating knowledge of extended temporal structure can be supported by replay and consolidation [33–36]; thus, updated representations may take time to emerge. For instance, during rest, the hippocampus reactivates sequences that were never directly experienced [33,34], and this reactivation may reorganize internal models in service of future behavior [37–39]. Fast and slow updating are not mutually exclusive: rapid initial updating may be followed by slower changes.

After assessing the time course of hippocampal updating, we asked *how* the hippocampus may integrate memories for initially separate, temporally extended sequences when these sequences become connected by a new link. We investigated two questions about how connected sequence representations were related to their pre-integration representations (as opposed to being newly-generated patterns not present pre-integration; [13]). First, we tested whether a newly integrated sequence representation preserved at least some features of the individual, original sequences, exhibiting relative *stability* even after updating. Second, we tested whether linking two sequences caused *blending* of their representations, such that the representation of one sequence inherited some of the original features of the other [1]. Stability and blending are not mutually exclusive: integration may result in representations that preserve elements of the original sequences, while also blending representations together or incorporating novel features.

To determine *when* and *how* hippocampal representations are updated after previously distinct sequences are connected, we asked participants to use sequence memories to anticipate events at multiple timescales (**Figure 1a**). After learning 4 sequences of virtual reality environments, participants learned novel, local event transitions that linked 2 of the previously separate sequences into a single, connected sequence (**Figure 1b**). The other 2 sequences remained separate and served as our baseline unconnected sequences. During fMRI, participants anticipated upcoming environments along both the connected and unconnected sequence conditions (**Figure 1c**). Critically, trials in the connected sequence required participants to traverse novel transitions. We examined patterns of activity in the hippocampus to: (1) determine whether trials in the connected vs. unconnected sequences were represented more similarly, consistent with integration; (2) establish the temporal dynamics by which integration occurred; (3) test if hippocampal integration supported prediction along the connected sequence; and (4) characterize whether updated hippocampal representations incorporated a novel, shared pattern that was not present for either sequence initially; retained stability with prior structure; and/or blended features of connected sequences.

**Figure 1.**
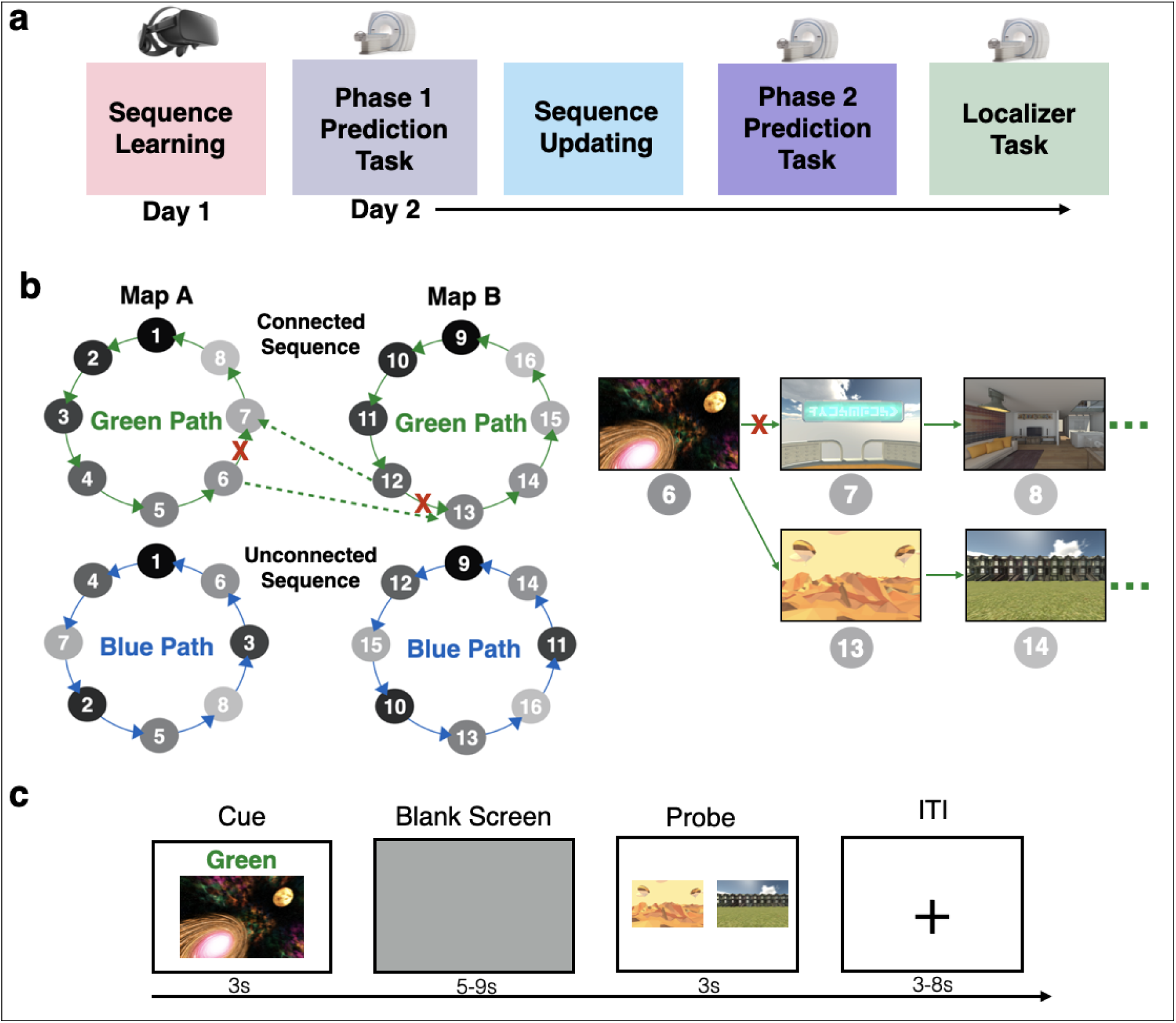
Sequence Structure and Prediction Task. **(a)** Experiment overview. Day 1 sequence learning took place during immersive virtual reality. Day 2 tasks took place during fMRI. **(b)** Sequence structure. Stimuli were 8 environments in Map A and a different 8 in Map B (gray nodes). Participants learned 4 sequences, 2 in each map. Each map had one green path and one blue path (solid lines), traversing the same 8 environments in a different order. Participants predicted upcoming environments across both maps and paths. They then learned that one environment in Map A was now connected to one in Map B (dashed lines), and vice versa, on either the green or blue path (counterbalanced). They were told the old connections between these “bridge” environments no longer worked (red X) and they could only use the new ones, forming a single, connected map for one path. The other path remained unchanged. To learn the new connection, participants viewed an environment from Map A (on either path) and were told it now connected to an environment in Map B, and vice versa. **(c)** Participants made anticipatory judgments. They were cued with an image from either the connected or unconnected sequence, along with a path cue (Green or Blue). A blank screen followed, during which they predicted upcoming environments in the sequence. They were then probed with 2 images of upcoming environments and had to choose which was coming up sooner in the cued sequence.

## Methods

### Dataset

We used data from Tarder-Stoll, Baldassano*, & Aly* [3]. 32 participants (21 female; age: 19-35 years, mean = 24.17, sd = 4.11; education: 13-29 years, mean = 16.85, sd = 3.76) learned sequences of environments in immersive virtual reality (VR) and then, during fMRI, anticipated upcoming environments (**Figure 1a**). All participants in the dataset gave written, informed consent in accordance with the Institutional Review Board at Columbia University. fMRI data are available on OpenNeuro: https://openneuro.org/datasets/ds005125. Data and code to reproduce analyses and figures in this paper are available at https://osf.io/pej92/.

### Task Overview

Sequence Learning and the Phase 1 Prediction Task are not of primary interest, but for completeness are briefly described here and described in full in Tarder-Stoll et al. [3]. In the Sequence Learning task, participants learned 2 sequences (“Green Path” and “Blue Path”) within each of 2 maps (Map A and Map B) in immersive VR, for a total of 4 sequences (**Figure 1b**). Map A and Map B contained 8 distinct environments each. Within each map, the Green Path and Blue Path contained the same environments in a different order. Participants learned the order of the 4 sequences by generating stories and then experiencing the environment sequences in immersive virtual reality using an Oculus Rift. Participants were then given a recall test to ensure they had learned all 4 sequences.

Participants returned one day later and completed the Phase 1 Prediction Task during fMRI (2 runs, 32 trials per run). In each trial, participants viewed an environment cue from a given map (A or B) and a path cue (Green or Blue) for 3 seconds. They then saw a blank screen for 5 to 9 seconds, followed by 2 images of upcoming environments. Participants indicated which of the 2 environments was coming up sooner in the sequence on the cued path (Green or Blue), relative to the cue environment. The correct answer could be 1 to 4 steps in the future. The incorrect answer was selected to be 1 to 4 steps in the future from the correct answer, with the constraint that it could be no more than 5 steps in the future from the cue. Participants had 3 seconds to respond, but were told that they could use the blank screen to predict upcoming environments in the order of the cued map and path. This task is analyzed in Tarder-Stoll et al., [3]. Here, this Phase 1 Prediction Task is solely used to explore how sequence representations change before vs. after sequence updating (see *fMRI Analysis)*.

#### Sequence Updating

Next, participants completed the Sequence Updating task. Participants were told that one environment in Map A was now connected to one environment in Map B on either the Green or Blue Path. One environment in Map B also connected back to Map A, creating a single connected sequence encompassing all environments in both maps (**Figure 1b)**. The path (Green or Blue) that was the connected sequence was counterbalanced across participants, with the other path remaining unconnected across maps and serving as the baseline condition. For example, if a participant learned that the Green Path now connected Map A and Map B, the two Blue Paths (in Map A and Map B) would serve as the unconnected paths.

For the connected sequence, the environment that linked Map A to Map B (a “bridge environment”) was randomly selected for each participant, as was the connecting environment in Map B. Following this new connection, the sequence of environments continued through all environments in Map B (in their previously learned order), and then linked back to Map A at the environment that had originally followed Map A’s bridge environment, forming a circle of all 16 environments (**Figure 1b**). To learn these new connections, participants viewed a video of the bridge environment in Map A transitioning to Map B on the connected sequence (Green or Blue). In the video, participants saw a panorama of the Map A bridge environment in virtual reality for ten seconds. After five seconds, a green and a blue sphere appeared, and a virtual hand touched one of the spheres (depending on whether the Green or Blue path was the connected sequence), triggering a transition to a panorama of the Map B bridge environment (see **Supplementary Materials** for an example). They then viewed a video of the bridge environment in Map B transitioning back to Map A. Participants were told to think about how the new connections could be incorporated into the stories that they had generated to learn the original sequences. Participants were then asked to recall the novel connections linking Map A and Map B. They were shown a screenshot of the Map A bridge environment and asked to verbally indicate which environment would come next, based on the connections they had just learned. After making their response, they were shown a screenshot of the correct next environment. Similarly, participants were also shown a screenshot of the Map B bridge environment, asked to verbally recall which environment would come next using the new connections, and subsequently shown a screenshot of the correct next environment. If participants responded incorrectly, they repeated the recall task until they could correctly indicate the next environment for both novel connections.

#### Phase 2 Prediction Task

Participants then completed the primary task of interest during fMRI – the Phase 2 Prediction Task using the updated sequences (**Figure 1c)**. Participants predicted upcoming environments on the connected and unconnected sequences (4 runs, 24 trials per run). Participants were cued with an environment from one map (A or B) along with a path cue (Green or Blue) for 3 seconds. Participants then viewed a blank screen for a variable duration (5 to 9 seconds). Then, participants were presented with 2 images of upcoming environments and judged which was coming up sooner in the cued sequence, relative to the cue image. Participants were given 3 seconds to respond. This relatively short response deadline was implemented to encourage participants to use the blank screen period to generate predictions along the cued path in preparation for the forced-choice decision. The correct answer could be 1 to 4 steps away from the cue image. The incorrect answer could be a maximum of 5 steps away. There was a uniformly sampled 3 to 8 second jittered inter-trial interval (ITI), during which participants viewed a fixation cross. At the end of each run, there was a 60 second rest period during which participants viewed a blank screen.

Critically, when cued with the connected sequence (Green or Blue Path, counterbalanced across participants), participants were told to use the new connections they just learned that linked Map A and Map B. Thus, participants could be cued with an environment from Map A and the correct answer could be in Map B (**Figure 1c**). On the unconnected sequences, Map A and Map B remained separate.

In each of the 4 runs (9 minutes each), participants were cued with every environment from Map A and Map B on the connected sequence (16 environments across both maps) and with half of the environments from Map A and B on the unconnected sequences (8 environments across both maps) for a total of 24 trials per run. In the probe phase, the correct answer was equally distributed across steps into the future (1 to 4). The incorrect answer was randomly selected with the constraints that it must be at least 1 step in the future from the correct answer but no more than 5 steps in the future from the cue. Within a run, participants completed trials from the connected sequence and unconnected sequence conditions, blocked by condition; block order was randomized across runs and participants. The order of the cues was randomized and intermixed across Map (A or B) within each block.

#### Localizer Task

Next, participants completed 4 runs of the Localizer Task (4 minutes each). This allowed us to select voxels that reliably discriminated between environments and obtain environment-specific template patterns of brain activity across those voxels (see *fMRI Analysis*). Participants viewed all environments in a randomized order. They were first shown an environment *without* a path cue (Green or Blue) for 1 second. The absence of the path cue allowed us to obtain a context-independent pattern of brain activity for each environment. Participants then saw a blank screen for 5 seconds, during which they were told to imagine themselves in the environment in immersive virtual reality, where the environments had been learned. Participants then saw 8 images of the environment, 45° apart, for 4 seconds, to mimic a panorama. Finally, participants were given 3 seconds to rate how vividly their imagination matched these images. There was a uniformly jittered ITI between 3 and 8 seconds.

### Behavioral Analysis

Behavioral data were analyzed in R with t-tests and generalized linear and linear mixed effects models (GLMMs and LMMs, glmer and lmer function in the *lme4* package; [40]). For analyses that modeled multiple observations per participant, such as accuracy or response time on a given trial, models included random intercepts and slopes for all within-participant effects. All response time models examined responses on correct trials only.

We first ensured that participants performed effectively during the Phase 2 Prediction Task, across both conditions, by comparing accuracy to chance (50%) using a one-sample t-test. We then conducted two one-sample t-tests comparing performance on the Phase 2 Prediction Task to chance separately for the connected and unconnected sequence conditions.

Next, we examined how performance changed across runs on the Phase 2 Prediction Task. We used a GLMM to model trial-wise accuracy as a function of condition (unconnected sequences = −0.5, connected sequence = 0.5), run (run 1 = −0.75, run 2 = −0.25, run 3 = 0.25, run 4 = 0.75), and their interaction.

Within the connected sequence condition, there were 3 trial types: (1) trials in which both probes were in the same map as the cue (same-map trials; these do not require crossing the bridge environment); (2) trials in which one probe was in a different map than the cue (one-different trials; these trials also do not require crossing the bridge environment because the environment in the same map is always the correct answer); and (3) trials in which both probes were in a different map than the cue (different-map trials; these trials require crossing the bridge environment; **Figure 2**). We tested whether accuracy differed as a function of trial type (same-map = 0, one-different = 1, different-map = 2), run (run 1 = −0.75, run 2 = −0.25, run 3 = 0.25, run 4 = 0.75), and their interaction. Trial type was a dummy-coded factor, allowing us to investigate how performance changed on one-different and different-map trials, relative to same-map baseline.

**Figure 2.**
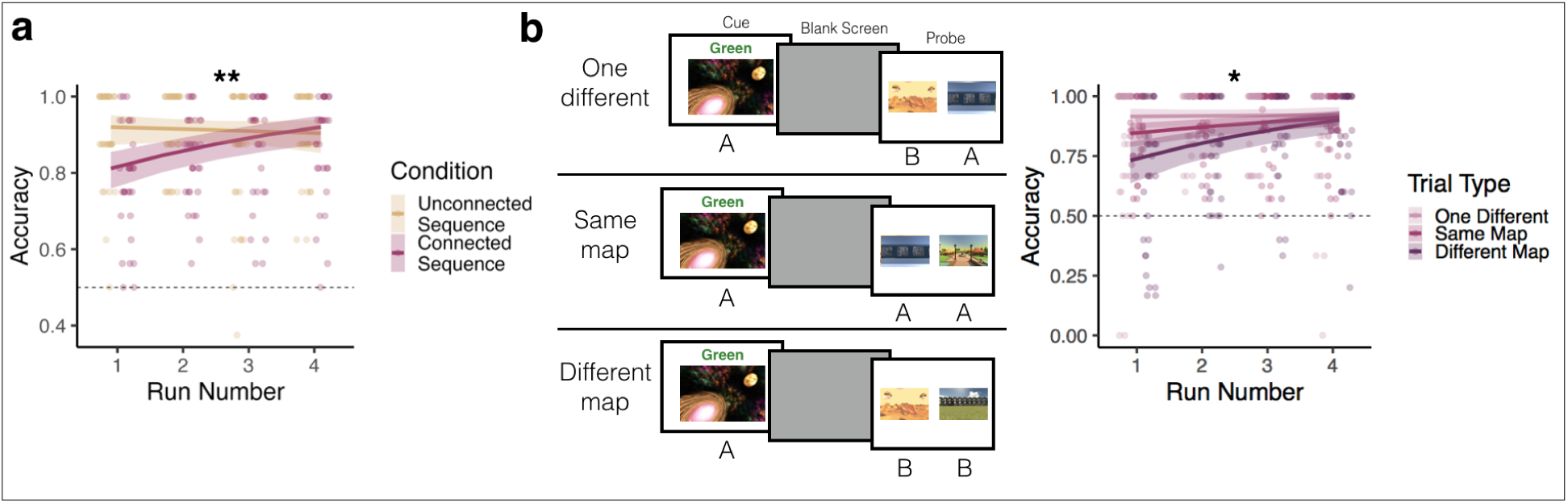
Prediction Task Performance. **(a)** Overall Behavioral Performance. Solid lines and error ribbons indicate model predictions with 95% confidence intervals; points indicate individual participants’ performance for each run. ** p < .001 for condition by run interaction. **(b)** Performance by trial type in the connected sequence condition. See main text for description of trial types. Performance was highest on one-different, followed by same-map, and then different-map trials. Performance improved the most across runs on different-map trials. Solid lines and error ribbons indicate model predictions with 95% confidence intervals; points indicate individual participants’ performance for each run. * p < 0.05 for trial type by run interaction.

### MRI Acquisition

MRI acquisition was reported in Tarder-Stoll et al. [3] and reiterated here for convenience. We collected whole-brain fMRI data using a 64-channel head coil on a 3 Tesla Siemens Magnetom Prisma scanner. We first acquired T1 structural scans using a magnetization-prepared rapid acquisition gradient-echo (MPRAGE) sequence. Functional images used a multiband echo-planar imaging (EPI) sequence (repetition time = 1.5s, echo time = 30ms, in-plane acceleration factor = 2, multiband acceleration factor = 3, voxel size = 2mm iso). There were 69 oblique axial slices collected in an interleaved order, with slices tilted −20 degrees relative to the AC-PC line. At the end of the session, we collected field maps (TR = 679 ms, TE = 4.92 ms/7.38 ms, flip angle = 60°, 69 slices, 2 mm isotropic).

### Preprocessing

Results included in this manuscript come from preprocessing performed using *fMRIPrep* 1.5.2 (Esteban, Markiewicz, et al. (2018); Esteban, Blair, et al. (2018); RRID:SCR_016216), which is based on *Nipype* 1.3.1 (Gorgolewski et al. (2011); Gorgolewski et al. (2018); RRID:SCR_002502).

#### Anatomical data preprocessing

The T1-weighted (T1w) image was corrected for intensity non-uniformity (INU) with N4BiasFieldCorrection (Tustison et al. 2010), distributed with ANTs 2.2.0 (Avants et al. 2008, RRID:SCR_004757), and used as T1w-reference throughout the workflow. The T1w-reference was then skull-stripped with a *Nipype* implementation of the antsBrainExtraction.sh workflow (from ANTs), using OASIS30ANTs as target template. Brain tissue segmentation of cerebrospinal fluid (CSF), white-matter (WM) and gray-matter (GM) was performed on the brain-extracted T1w using fast (FSL 5.0.9, RRID:SCR_002823, Zhang, Brady, and Smith 2001). Brain surfaces were reconstructed using recon-all (FreeSurfer 6.0.1, RRID:SCR_001847, Dale, Fischl, and Sereno 1999) [81], and the brain mask estimated previously was refined with a custom variation of the method to reconcile ANTs-derived and FreeSurfer-derived segmentations of the cortical gray-matter of Mindboggle (RRID:SCR_002438, Klein et al. 2017). Volume-based spatial normalization to one standard space (MNI152NLin2009cAsym) was performed through nonlinear registration with antsRegistration (ANTs 2.2.0), using brain-extracted versions of both T1w reference and the T1w template. The following template was selected for spatial normalization: *ICBM 152 Nonlinear Asymmetrical template version 2009c* [Fonov et al. (2009) [83], RRID:SCR_008796; TemplateFlow ID: MNI152NLin2009cAsym].

#### Functional data preprocessing

For each of the 10 BOLD runs found per subject (across all tasks and sessions), the following preprocessing was performed. First, a reference volume and its skull-stripped version were generated using a custom methodology of *fMRIPrep*. A deformation field to correct for susceptibility distortions was estimated based on a field map that was co-registered to the BOLD reference, using a custom workflow of *fMRIPrep* derived from D. Greve’s epidewarp.fsl script and further improvements of HCP Pipelines (Glasser et al. 2013). Based on the estimated susceptibility distortion, an unwarped BOLD reference was calculated for a more accurate co-registration with the anatomical reference. The BOLD reference was then co-registered to the T1w reference using bbregister (FreeSurfer) which implements boundary-based registration (Greve and Fischl 2009). Co-registration was configured with six degrees of freedom. Head-motion parameters with respect to the BOLD reference (transformation matrices, and six corresponding rotation and translation parameters) are estimated before any spatiotemporal filtering using mcflirt (FSL 5.0.9, Jenkinson et al. 2002). The BOLD time-series, were resampled to surfaces on the following spaces: *fsaverage6*. The BOLD time-series (including slice-timing correction when applied) were resampled onto their original, native space by applying a single, composite transform to correct for head-motion and susceptibility distortions. These resampled BOLD time-series will be referred to as *preprocessed BOLD in original space*, or just *preprocessed BOLD*. The BOLD time-series were resampled into standard space, generating a *preprocessed BOLD run in [‘MNI152NLin2009cAsym’] space*. First, a reference volume and its skull-stripped version were generated using a custom methodology of *fMRIPrep*. Several confounding time-series were calculated based on the *preprocessed BOLD*: framewise displacement (FD), DVARS and three region-wise global signals. FD and DVARS are calculated for each functional run, both using their implementations in *Nipype* (following the definitions by Power et al. 2014). The three global signals are extracted within the CSF, the WM, and the whole-brain masks. Additionally, a set of physiological regressors were extracted to allow for component-based noise correction (*CompCor*, Behzadi et al. 2007). Principal components are estimated after high-pass filtering the *preprocessed BOLD* time-series (using a discrete cosine filter with 128s cut-off) for the two *CompCor* variants: temporal (tCompCor) and anatomical (aCompCor). tCompCor components are then calculated from the top 5% variable voxels within a mask covering the subcortical regions. This subcortical mask is obtained by heavily eroding the brain mask, which ensures it does not include cortical GM regions. For aCompCor, components are calculated within the intersection of the aforementioned mask and the union of CSF and WM masks calculated in T1w space, after their projection to the native space of each functional run (using the inverse BOLD-to-T1w transformation). Components are also calculated separately within the WM and CSF masks. For each CompCor decomposition, the *k* components with the largest singular values are retained, such that the retained components’ time series are sufficient to explain 50 percent of variance across the nuisance mask (CSF, WM, combined, or temporal). The remaining components are dropped from consideration. The head-motion estimates calculated in the correction step were also placed within the corresponding confounds file. The confound time series derived from head motion estimates and global signals were expanded with the inclusion of temporal derivatives and quadratic terms for each (Satterthwaite et al. 2013). Frames that exceeded a threshold of 0.5 mm FD or 1.5 standardised DVARS were annotated as motion outliers. All resamplings can be performed with *a single interpolation step* by composing all the pertinent transformations (i.e. head-motion transform matrices, susceptibility distortion correction when available, and co-registrations to anatomical and output spaces). Gridded (volumetric) resamplings were performed using antsApplyTransforms (ANTs), configured with Lanczos interpolation to minimize the smoothing effects of other kernels (Lanczos 1964). Non-gridded (surface) resamplings were performed using mri_vol2surf (FreeSurfer).

Many internal operations of *fMRIPrep* use *Nilearn* 0.5.2 (Abraham et al. 2014, RRID:SCR_001362), mostly within the functional processing workflow. For more details of the pipeline, see the section corresponding to workflows in *fMRIPrep*’s documentation (https://fmriprep.org/en/latest/workflows.html).

#### Copyright Waiver

The above boilerplate text was automatically generated by fMRIPrep with the express intention that users should copy and paste this text into their manuscripts *unchanged*. It is released under the CC0 license.

### fMRI Analysis

Analyses were performed in Python and R. Representational similarity analyses [41] were performed using custom code in Python 3. Statistical analysis comparing pattern similarity across conditions, correlations between fMRI results and behavior, and visualizations were performed using custom code in R.

#### Conjunction ROI Definition

Our primary region of interest (ROI) was the hippocampus. We also conducted analyses in early visual cortex as a control ROI. We used a conjunction approach to identify voxels within these regions that reliably discriminated between environments (as in [3]). We defined an anatomical hippocampus ROI from the Harvard-Oxford probabilistic atlas in FSL (threshold: p = 0.50) and a visual cortex ROI (V1-V4) from the probabilistic human visual cortex atlas provided in Wang et al ([42]; threshold: p = 0.50). The ROIs and functional data were both transformed into 2 mm MNI space (MNI152NLin2009cAsym). We then identified voxels within these ROIs whose responses to the environments were consistent across participants using the same approach from Tarder-Stoll et al. [3], which was adapted from Tarhan and Konkle [43].

We first conducted a whole-brain GLM (using custom code in Python) predicting univariate activity from task and nuisance regressors during the Localizer Task. For each participant, we modeled BOLD activity for each of the 16 environments, collapsed across runs. The regressors for each environment included both viewing and imagination periods (see *Methods/Task Overview*). The model also included nuisance regressors: translation and rotation along the X, Y, and Z axes and their derivatives, motion outliers as determined by fMRIprep, CSF, white matter, framewise displacement, and discrete cosine-basis regressors for periods up to 125 seconds). We then extracted the beta weights for each environment for each participant from the whole-brain GLM for each voxel in our atlas-based hippocampus and visual cortex ROIs. We then computed, for each voxel, the Pearson correlation between each participant’s vector of beta weights (one beta weight for each environment) and the average vector of the remaining participants. An environment reliability score was calculated by averaging these correlations across all iterations of the held-out participant, and we only included voxels with an environment reliability score of 0.1 or greater in our final analyses. Thus, ROI selection was conducted using a set of data that was independent from the main analyses of the Prediction Task. We used these conjunction ROIs (156 voxels in hippocampus; 2931 voxels in visual cortex) for subsequent analyses.

#### Phase 2 Prediction Task Analysis

We used custom code in Python to run GLMs predicting whole-brain univariate BOLD activity from task and nuisance regressors from the Phase 2 Prediction Task. For each participant, we modeled BOLD activity separately for each run with regressors for the cue, blank screen, and probe periods, separately for each cued environment in Map A and Map B (1 to 16) and for each path (Green Path and Blue Path). This resulted in 32 task regressors for each phase (cue, blank screen, probe) of the Phase 2 Prediction Task per run. These task regressors were convolved with the one-parameter gamma hemodynamic response function from AFNI’s 3dDeconvolve. We also included nuisance regressors (translation and rotation along the X, Y, and Z axes and their derivatives, motion outliers as determined by fmriprep, CSF, white matter, framewise displacement, and discrete cosine-basis regressors for periods up to 125 seconds). The resulting beta weights for our task regressors for each run were examined within our hippocampal and visual cortex conjunction ROIs. The same GLM approach (with the same task and nuisance regressors) was used to model the Phase 1 Prediction Task, for the purposes of comparing Phase 1 and Phase 2 activity patterns (see Results).

Pattern similarity analyses were conducted to examine how sequence representations in hippocampus and visual cortex changed across maps in the connected vs. unconnected sequence conditions. For each participant, we obtained the correlation between (1) the blank screen pattern of activity on each trial and (2) the blank screen pattern of activity for all other trials in the same condition (connected sequence or unconnected sequences) but with a cue from the other map (A or B) (**Figure 3a**). For example, if the cue environment on a given trial was from Map A and in the connected sequence, we obtained the correlation between that trial’s blank screen activity pattern and the blank screen activity patterns for trials in which the cue environment was in Map B and in the connected sequence. To explore how across-map pattern similarity changes across the Phase 2 Prediction Task while accounting for temporal autocorrelation within a run, we conducted this analysis across neighboring runs separately for early runs (1 & 2) and late runs (3 & 4). For example, if a trial was from run 1, we obtained the correlation with trials from the same condition and different map in run 2 and vice versa (**Figure 3a**).

**Figure 3.**
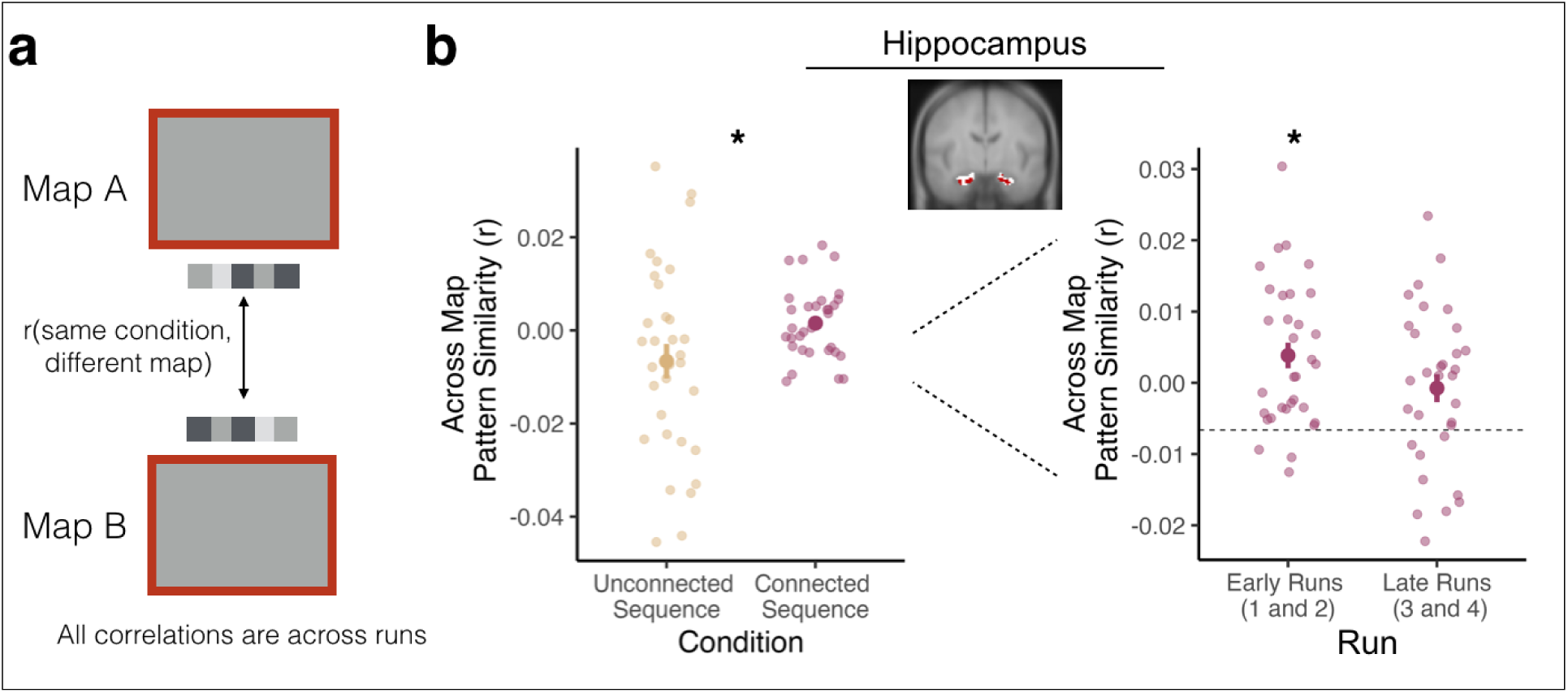
Sequence Integration in Hippocampus. **(a)** Pattern similarity analysis. To quantify across-map pattern similarity during the Phase 2 Prediction Task, we obtained the correlation between (1) the blank screen pattern of activity on each trial and (2) the blank screen pattern of activity for all other trials in the same condition (connected sequence or unconnected sequences) but from the different map (A or B) for each participant. **(b)** Pattern similarity results. Left: We investigated across-map pattern similarity within our hippocampus conjunction ROI, collapsed across early and late runs. Across-map pattern similarity was higher for the connected sequence compared to the unconnected sequences. Note that the higher variance in the unconnected sequence condition is likely due to fewer trials in this condition. Right: Across-map pattern similarity was higher in early runs of the connected sequence condition compared to the unconnected sequence condition. There was no significant difference between conditions in late runs. Large points indicate the group average across-map pattern similarity, error bars indicate standard error of the mean, and small, transparent points indicate each participant’s across-map pattern similarity. Dashed line indicates group average value in the unconnected sequence. * p < .05

The resulting correlations indicate the similarity between patterns of activity for Map A and Map B across runs. We tested for significant differences between the connected and unconnected sequence conditions with a paired-samples t-test. We ran follow-up paired-sample t-tests to assess differences in pattern similarity for early and late runs in the connected sequence, compared to averaged pattern similarity across all runs in the unconnected sequence condition. We opted to average across runs in the unconnected sequence condition because (1) we did not find differences in pattern similarity across runs for this condition and (2) we did not have enough trials to reliably estimate run-wise pattern similarity values in the unconnected sequences (8 trials per run, half the number of trials as the connected sequence; [44]). These analyses, which separately compare the first two runs and the last two runs of the connected sequence condition to all four runs of the unconnected sequence condition, match the number of trials used for each condition in each comparison. This analysis therefore addresses the imbalanced trial numbers in the analysis across all runs, because the connected sequence condition had double the number of trials as the unconnected sequence across all four runs.

To determine if pattern similarity was driven by changes in overall activity, we conducted univariate analyses by obtaining the average beta weight across all voxels in our hippocampus conjunction ROI, separately for the connected and unconnected sequence conditions. We conducted these analyses across all runs, and separately for early and late runs. Further, we separately examined Cue and Blank Screen periods of the Phase 2 Prediction Task (**Figure 1c**). For each approach we compared univariate activity in the connected and unconnected sequence conditions using paired sample t-tests.

#### Relationship Between Prediction Task Pattern Similarity and Behavior

We determined whether across-map pattern similarity in hippocampus or visual cortex was related to performance (**Figure 4**). We performed an individual differences analysis in which we used the unconnected sequences condition as a baseline and examined whether changes in neural representations for the connected vs. unconnected sequences were related to behavioral differences across these conditions. We obtained the Spearman rank-order correlation between (1) the difference between participants’ accuracy in the connected vs. unconnected sequence condition and (2) the difference between participants’ hippocampal across-map pattern similarity in the connected vs. unconnected sequence condition.

**Figure 4.**
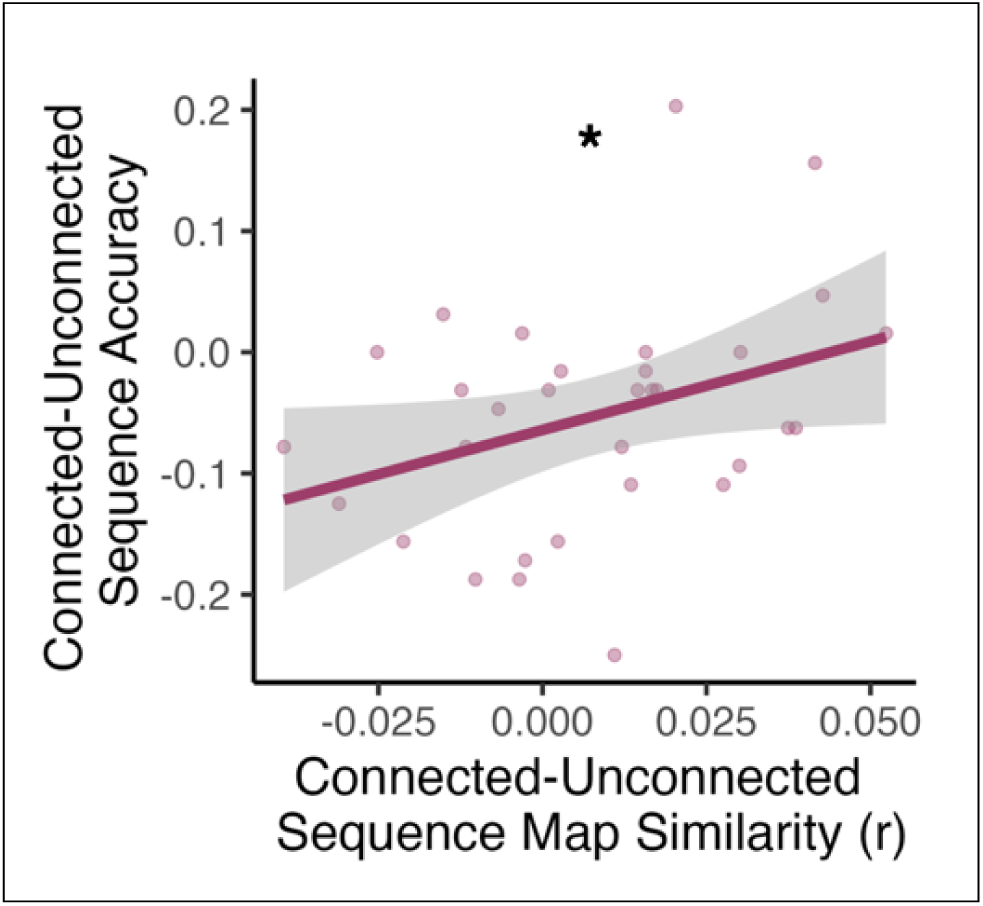
Across-map pattern similarity in the hippocampus supports behavioral predictions. Greater across-map pattern similarity in the hippocampus for connected vs. unconnected sequences was associated with reduced behavioral costs for the connected sequence during the Prediction Task. Pink line and gray error ribbons indicate the correlation with 95% confidence intervals; points indicate each participant’s across-map pattern similarity and Prediction Task performance. * p < .05

#### Tests of Representational Stability, Blending, and Novel Patterns

We conducted three analyses to investigate the nature of updated representations in the hippocampus for the connected sequences. One analysis examined if there was *stability* of representations for a given sequence from pre- to post-updating. The second analysis looked for *blending* of sequence representations after updating. The third analysis looked for evidence of a new, shared representation that was not present in either sequence initially (**Figure 5**).

**Figure 5.**
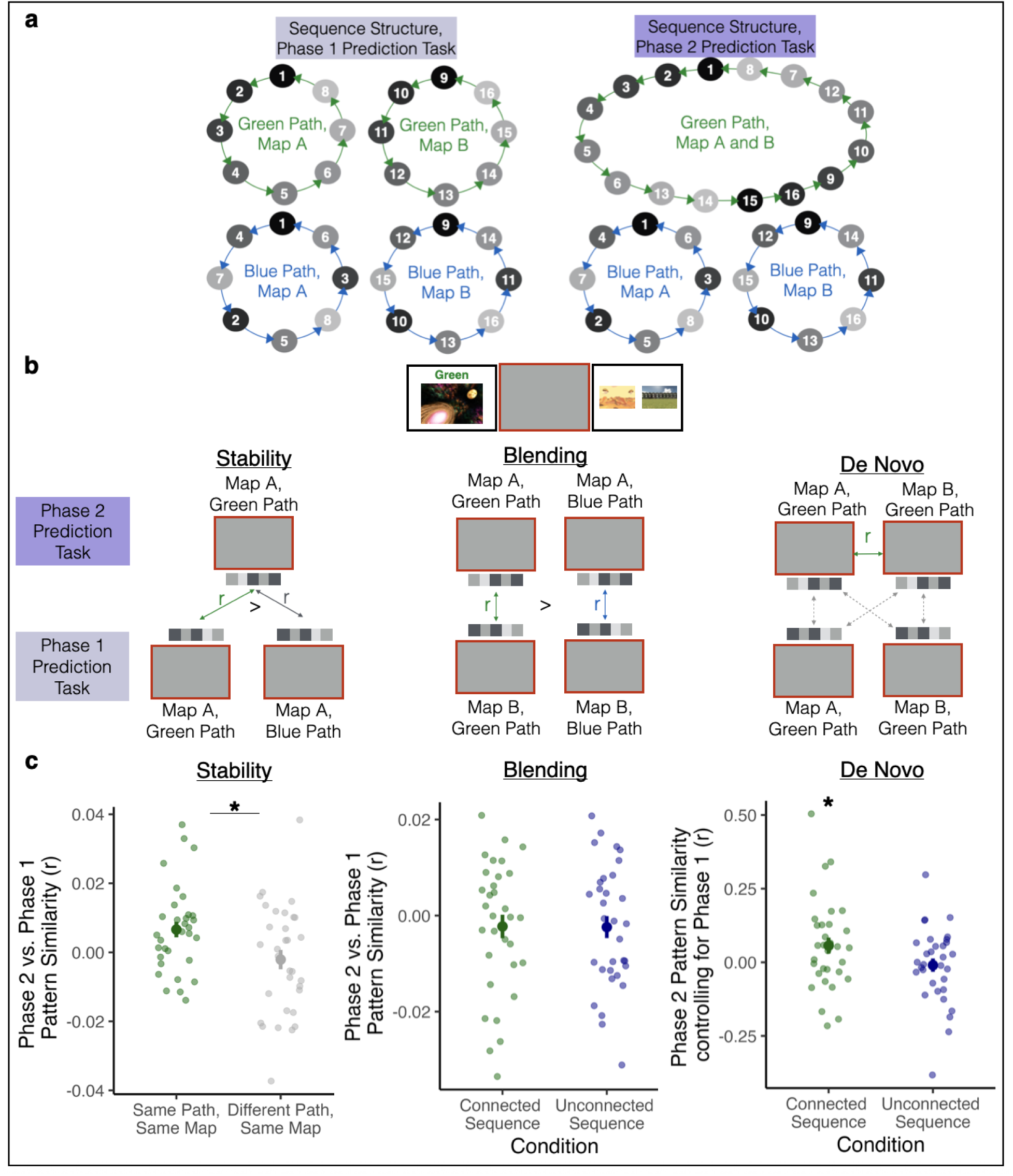
Stability and Blending of Hippocampal Representations. **(a)** Sequence structure in the Phase 1 Prediction Task and the Phase 2 Prediction Task. In this example, the green path is the connected sequence and the blue path is the unconnected sequence. **(b)** We analyzed patterns of brain activity during the blank screen period of the Phase 1 and Phase 2 Prediction Task to investigate stability, blending, and de novo representations in the connected vs. the unconnected sequences. To look for stability in representations, we obtained the correlation between the Phase 2 Prediction Task activity patterns for Map A Green Path trials (connected sequence) and (1) the Phase 1 Prediction Task activity patterns for Map A Green Path trials (same sequence, pre-updating: “same path, same map”) and (2) Phase 1 Prediction Task activity patterns for Map A Blue Path trials (unconnected sequence, “different path, same map”). We completed the same correlations for Map B as well (not shown). Higher correlations between the same path vs. different path would provide evidence of stable representations after integration. To look for evidence of blending, we obtained the correlation between the Phase 2 Prediction Task patterns of activity for Map A Green Path trials (connected sequence) and the Phase 1 Prediction Task patterns of activity for Map B Green Path trials (connected sequence), and vice versa. We then conducted the same comparisons for the Blue Path (unconnected sequence). Higher correlations for the connected vs. unconnected sequence would provide evidence for blended representations after integration. To look for evidence of a de novo representation, we obtained the partial correlation between the Phase 2 patterns of Map A and Map B in the connected sequence condition, after controlling for each of the Phase 1 patterns in that condition (dotted gray arrows). We conducted the same partial correlation analysis for the Blue Path (unconnected sequence). Partial correlations greater than 0 provide evidence for a de novo representation. **(c)** Left: Phase 2 vs. Phase 1 pattern similarity was higher for the same path vs. different path, indicating stable representations after updating. Middle: Across-map Phase 2 vs. Phase 1 pattern similarity was not significantly different for the connected sequence vs. the unconnected sequence, indicating no evidence for blended representations. Right: The partial correlation between Map A and Map B in the connected sequence was statistically significant after controlling for each of these representations’ similarity to pre-integration patterns. The partial correlation in the unconnected sequence was not statistically significant. Large points indicate the group average pattern similarity, error bars indicate standard error of the mean, and small, transparent points indicate each participant’s across-map pattern similarity. * p < .05

To investigate if there was *stability* of individual sequence representations in the newly connected sequence, we tested if trials from a given sequence after sequence updating were more similar to trials in the same sequence before updating than they were to trials in the same map for the unconnected sequence condition. Specifically, for each participant, we obtained the correlation between the blank screen activity pattern during the Phase 2 Prediction Task for each connected sequence trial and two types of trials from the Phase 1 Prediction Task: (1) trials from the same map in the connected sequence, and (2) trials from the same map in the unconnected sequence. For example, if a Phase 2 Prediction Task trial came from Map A in the connected sequence, we calculated its similarity to Phase 1 Prediction Task trials from Map A in the connected sequence (same sequence) and from Map A in the unconnected sequence (same environments in a different order). We compared these pattern similarity values using a paired-samples t-test. This allowed us to assess whether Phase 2 representations in the connected sequence were more similar to their own Phase 1 representations than to the Phase 1 representations of the same map in the unconnected sequence condition. Critically, because Map A (and, separately, Map B) in the connected and unconnected sequence conditions have the same environments in a different order, this comparison allows us to control for pattern similarity driven only by repetition of the same environments.

We next tested whether updated representations in the connected sequence condition reflected a *blend* of Map A and Map B sequences. If so, post-updating representations of Maps A and B in the connected sequence may “inherit” features from the newly connected sequence’s representations prior to learning. That is, Map A post-updating may contain features of Map B from pre-updating and vice versa. To test this, for each participant, for both the connected and unconnected sequences, we obtained the correlation between (1) the blank screen pattern of activity on each trial in the Phase 2 Prediction Task and (2) the blank screen pattern of activity for all trials with a cue from the other map (A or B) in the Phase 1 Prediction Task. For example, if the cue environment on a Phase 2 Prediction Task trial came from Map A, we obtained the correlation between its blank screen activity pattern and activity patterns from Phase 1 Prediction Task trials with cue environments from Map B. This was done for the connected sequence and unconnected sequences separately, and those two values were then compared with a paired samples t-test. This allowed us to determine if representations carried over from Phase 1 to Phase 2 more strongly for the connected vs. unconnected sequences.

Finally, to look for evidence of a *novel post-integration pattern* that was not present in either pre-integration patterns separately, we conducted a partial correlation analysis that examined similarity between Phase 2 patterns after controlling for the presence of each Phase 1 pattern in the same condition. For example, if the Green Path was connected for a given participant, we examined the partial correlation between the Phase 2 patterns for the Green Path in Map A and the Green Path in Map B, after regressing out any variance in these Phase 2 patterns related to the Phase 1 patterns for the Green Path in Map A and the Green Path in Map B. The same approach was used for the unconnected sequence condition as a control. This analysis allowed us to look for evidence that there was a new, shared pattern in the representations of the connected sequences after integration.

## Results

### Prediction Task Performance

In the Phase 2 Prediction Task, participants chose the probe that was coming up sooner relative to the cued environment 85.97% of the time, which was significantly above chance (50%, t(31) = 51.057, p < 0.000001). Performance was high and significantly above chance in both the unconnected sequence condition (M_accuracy_ = 89.453%, t(31) = 59.774, p < 0.000001) and the connected sequence condition (M_accuracy_ = 84.228%, t(31) = 41.814, p < 0.000001).

To determine how sequence memories are updated with experience, we tested how performance on the Phase 2 Prediction Task varied by condition and run. Performance was significantly higher in the unconnected vs. connected sequence condition (beta = −0.389, 95% CI = [-0.741, −0.037], p = 0.03; **Figure 2a**). Overall, performance did not significantly differ across runs of the Phase 2 Prediction Task (beta = 0.244, 95% CI = [-0.024, 0.513], p = 0.074; **Figure 2a**). However, there was a significant interaction between condition and run (beta = 0.742, 95% CI = [0.209, 1.276], p = 0.006; **Figure 2a**), such that performance increased across runs in the connected sequence condition (beta = 0.62, 95% CI = [0.329, 0.911], p = 0.00003), but not the unconnected sequence condition (beta = −0.122, 95% CI = [-0.563, 0.320], p = 0.588). In fact, performance in the connected sequence condition reached that of the unconnected sequence condition by the end of the task (beta = −0.150, 95% CI = [-0.874, 0.573], p = 0.684). Strikingly, even in the first run of the Phase 2 Prediction Task, performance was high and significantly above chance in the connected sequence condition (average accuracy = 78.52%, t(31) = 30.538, p < 0.000001), suggesting that participants rapidly updated their memories to reflect the connected sequence structure.

We next examined how performance changed as a function of run and trial type in the connected sequence condition. Trials in this condition differed in difficulty based on whether participants needed to mentally traverse between maps to correctly anticipate upcoming environments (**Figure 2b**, see *Methods*). Trials in which both probes were in the same map as the cue (same-map trials) should be easier than trials in which both probes were in the different map from the cue (different-map trials), because different-map trials required participants to cross the bridge environment to correctly respond. Trials in which only one of the probes is in the different map from the cue (one-different trials) should be easier than both same-map and different-map trials, because a participant should be able to reject the incorrect probe on the basis of map identity alone, rather than temporal order. Specifically, because the correct answer was at most four steps away, it is never possible for the different-map environment to be closer than the same-map environment. Because neither same-map nor one-different trials required crossing the bridge environment, the correct probe is the same for the updated sequence as it was for the original sequence. However, for different-map trials, neither option in the probe display was present in the original sequence and thus these trials required using the new connections to make a correct response. The different-map probes therefore provide an implicit feedback signal if a participant fails to use the updated connection between maps, because in this case neither of the probes would match their predictions; this indicates to the participant that they failed to follow the new post-bridge connections, and may prompt more rehearsal of those connections to support improved performance over time.

Indeed, accuracy in the connected sequence condition was highest on one-different trials (mean = 88.9%), followed by same-map trials (mean = 87.8%), and lowest on different-map trials (mean = 78.5%). There was a trend toward lower accuracy on different-map compared to same-map trials (beta = −0.368, 95% CI = [-0.744, 0.007], p = 0.054; **Figure 2b**), but there was no difference between one-different and same-map trials (beta = 0.284, 95% CI = [-0.139, 0.708], p = 0.188). Critically, there was a trial type by run interaction (beta = 0.585, 95% CI = [0.078, 1.091], p = 0.024; **Figure 2b**), such that performance on different-map trials increased across runs more than performance on the baseline same-map trials. In the first run of the Phase 2 Prediction Task, accuracy was significantly lower on different-map trials compared to same-map trials (beta = −0.8726, 95% CI = [-1.361, −0.384], p = 0.0005), but performance on different-map trials reached that of same-map trials in the final run (beta = −0.086, 95% CI = [-0.701, 0.529], p = 0.784). Importantly, performance on different-map trials was above chance even in the first run (t(31) = 3.340, p = 0.002), again showing successful rapid updating of predictions in the connected sequence. In contrast, there was no interaction between same-map and one-different trials across runs (beta = −0.002, 95% CI = [-0.650, 0.645], p = 0.994). Together, these analyses show that trials in which participants had to mentally traverse the bridge environments linking Map A and B (i.e., different-map trials) improved the most over the course of the Phase 2 Prediction Task.

Thus, participants updated their memories of temporal structure after a single learning episode to successfully make predictions. Sequence memories reflected the new sequence structure even in the first run of the Phase 2 Prediction Task, but behavior continued to improve – implying both rapid and gradual updating of learned representations.

### Integration of Hippocampal Sequence Representations

We next explored how updated sequences are represented in the hippocampus. We used multivoxel pattern analysis to determine if the connected sequence was represented more similarly than the unconnected sequences, consistent with hippocampal integration mechanisms [12]. We obtained the correlation between multivoxel patterns of activity elicited during the blank screen of the Phase 2 Prediction Task for trials in which cue environments came from Map A vs. Map B, separately for the connected and unconnected sequence conditions (see *Methods*; **Figure 3a**). Across-map pattern similarity was higher in the connected vs. unconnected sequence condition (t(31) = 2.070, p = 0.041; **Figure 3b**), indicating that hippocampal representations of Map A and Map B sequences were more similar when they were connected by the bridge environments. This confirms that the hippocampus supports updating of sequence memories by forming an integrated representation of the previously separate sequences. In contrast, across-map pattern similarity in visual cortex did not differ between the connected vs. unconnected sequence conditions (t(31) = −0.279, p = 0.781). Likewise, a follow-up exploratory analysis on regions of interest in retrosplenial cortex and the parahippocampal place area [3,45] revealed that neither of these regions showed a difference in across-map similarity across conditions (PPA: t(31) = −0.764, P = 0.45); RSC: t(31) = −0.154, p = 0.878).

We then asked how hippocampal across-map similarity varied as a function of the distance between a cue environment and the bridge environment to the other map. Participants did not know during the blank screen period how far away the probes would be, but the closest probe environment on each trial was at most 4 steps away from the cue; therefore only cue environments that could reach the other map within 4 steps could potentially require across-map integration to correctly select the closest probe. To determine if this had an impact on hippocampal patterns, we compared across-map pattern similarity in the connected sequence for environments that could reach the other map within 4 steps vs. could not reach the other map within 4 steps. We compared these two types of across-map pattern similarity for the connected sequence condition to the across-map pattern similarity of all environments in the unconnected sequence condition.

Across-map pattern similarity for the connected sequence was higher than that of the unconnected sequence condition for environments that could reach the other map within 4 steps (t(31) = 2.247, p = 0.026). The connected vs. unconnected sequence comparison was not significantly different for environments that could not reach the other map within 4 steps (t(31) = 0.748, p = 0.455). However, across-map pattern similarity was not significantly different for environments that were within vs. not within 4 steps of the other map (t(31) = 1.284, p = 0.406). This result suggests that across-map integration in the hippocampus may be elevated for those environments that are closest to the bridge; however, the lack of a significant difference between environments closer to vs. farther from the bridge indicates that future work is needed to establish the robustness of this effect.

To probe how hippocampal sequence representations are updated over time, we separately compared across-map pattern similarity in the connected sequence for early and late runs to the averaged across-map pattern similarity in the unconnected sequences. In early runs, hippocampal across-map pattern similarity was significantly higher in the connected vs. unconnected sequence condition (t(31) = 2.665, p = 0.012; **Figure 3b**). In late runs, across-map pattern similarity was not different in the connected vs. unconnected sequence conditions (t(31) = 1.304, p = 0.203; **Figure 3b**). There was a trending difference between early vs. late run across-map pattern similarity in the connected sequence condition (t(31) = −1.78, p = 0.085). We address this pattern in the Discussion. We then did the same early vs. late run analysis in visual cortex, and there was no difference in across-map pattern similarity between conditions, neither in early runs (t(31) = 0.244, p = 0.808) nor late runs (t(31) = 0.342, p = 0.734). Together, these results show that the hippocampus, but not visual cortex, integrates sequence representations in light of new information, and that this integration in hippocampus appears rapidly – in the first runs after new learning.

Integration of associated stimuli is linked to overall activity in the hippocampus [46]. We therefore investigated whether representational differences between conditions were due to differences in hippocampal univariate activity. Unlike across-map pattern similarity, there were no differences in hippocampal univariate activity between the connected and unconnected sequence conditions, either during the cue (t(31) = −1.404, p = 0.170) or the blank screen (t(31) = −0.081, p = 0.936) portion of the Phase 2 Prediction Task. There was no significant difference between conditions during early runs (cue: t(31) = −1.105, p = 0.278; blank screen: t(31) = −0.710, p = 0.483) or during late runs (cue = t(31) = −0.871, p = 0.390; blank screen: t(31) = 0.476, p = 0.638). Thus, the increase in across-map pattern similarity in the connected vs. unconnected sequence condition cannot be attributed to overall changes in hippocampal univariate activity.

### Rapid Sequence Integration Supports Memory Updating

We next explored whether rapid sequence updating in the hippocampus was related to behavior. Overall, participants tended to show behavioral costs for the connected vs. unconnected sequence condition, at least in early runs (**Figure 2a**). We asked whether participants who showed more evidence for hippocampal integration for the connected sequence, compared to the baseline unconnected sequences, exhibited reduced behavioral costs (or behavioral benefits) for the connected vs. unconnected sequences. We obtained the Spearman’s rank-order correlation between (1) the difference in hippocampal across-map similarity in the connected vs. unconnected sequence conditions and (2) the behavioral difference between the connected vs. unconnected sequence conditions (see *Methods*). As a control analysis, we also calculated the Spearman’s rank-order correlation with across-map pattern similarity in visual cortex. We found a positive relationship in the hippocampus (Spearman’s *rho* = 0.368, p = 0.038; **Figure 4a**): participants who showed the most across-map integration for the connected (vs. unconnected) sequence exhibited reduced behavioral costs (or even behavioral benefits) in the connected (vs. unconnected) sequence condition. The same analysis in visual cortex was not statistically significant (Spearman’s *rho* = 0.055, p = 0.763). Thus, increased across-map pattern similarity in hippocampus, but not visual cortex, was associated with reduced costs to prediction performance after sequence updating.

### Novel Hippocampal Representations After Sequence Updating

What is the nature of the post-updating, integrated representations in the hippocampus? Previously separate sequences become more similar after new learning (**Figure 3**), which could be due to the creation of a *new (shared) representation*, whose features were not present in either sequence prior to integration [13]. It is also possible that some features of the pre-integration representations persist even after sequence updating. For example, the two connected sequences may maintain some *stability* with their individual pre-updating representations. Another (not mutually exclusive) possibility is that across-map integration results in a mixture of the two original sequences — with *blended* representations of Map A and Map B in the connected, but not the unconnected, sequence. To explore these possibilities, we conducted three analyses to determine if hippocampal patterns in the connected sequence showed evidence for stability, blending, or the creation of a novel, shared representation.

To look for *stability in representations*, we tested whether each component of the connected sequence (Map A and Map B environments in the connected path) showed similarity with their *pre-updating* representations (**Figure 5**). We compared these values to a baseline that controlled for pattern similarity driven by repetition of the same visual images, by obtaining the correlation between the Phase 2 representations of each Map in the connected sequence and the Phase 1 representations of the same Map in the unconnected sequence condition, which had the same environments in a different order. For example, the stability of the connected sequence in Map A was assessed by determining if its Phase 2 representation was more similar to its Phase 1 representation than to the Phase 1 representation of the unconnected sequence in Map A. Thus, this analysis allowed us to examine whether sequences that became connected retained similarity to their original representations, above and beyond any generic pattern similarity driven by repetition of the same cue images (although the images are no longer on the screen during the blank period that was analyzed, cognitive processes evoked by scene perception are likely to linger during the blank period; subtraction of baseline pattern similarity for the same images therefore allows us to better isolate effects related to anticipation of upcoming images in the cued sequence, rather than sequence-independent processing of the cue image).

We found evidence for stability in hippocampal representations for the connected sequences, such that their post-updating representations retained features of their pre-updating representations (t(31) = 2.15, p = 0.039). Thus, the hippocampus can rapidly update representations of sequences in a way that partially maintains their prior identities but also links them together.

To look for evidence of *blended representations* in the connected sequences, we tested whether Map A after updating became more similar to Map B before updating (and vice versa) in the connected sequence condition compared to the same comparisons for the unconnected sequence condition. This would suggest that after sequences became connected, Map A inherited features of Map B, and vice versa – beyond the level of blending present for the unconnected sequences. We found no evidence for blended representations of the two sequences after updating (t(31) = 0.037, p = 0.971). Map A after updating was not more similar to Map B before updating (and vice versa) in the connected sequence condition relative to the same comparison in the unconnected sequence condition.

Finally, we tested whether Map A and Map B in the connected sequence condition showed evidence for a *novel, shared representation* that was not present in either representation prior to integration. To do so, we examined the partial correlation between the Phase 2 patterns of Map A and Map B in the connected sequence condition, after controlling for each of the Phase 1 patterns in that condition. The partial correlation between Map A and Map B in the connected sequence was statistically significant after controlling for each of these representations’ similarity to pre-integration patterns (t(31) = 2.09, p = 0.045), showing the creation of a new, shared pattern that was not present in either pattern pre-integration. This partial correlation was not statistically significant for the unconnected sequence condition (t(31) = −0.44, p = 0.67), and the difference between conditions was marginally significant (t(31) = 1.8965, p = 0.063). Together, these analyses show that the enhanced post-updating similarity we observed (**Figure 3b**) is at least partly driven by the incorporation of a novel pattern (not present pre-integration) into both Maps of the connected sequence.

## Discussion

We investigated when and how temporally extended sequences become integrated in the brain to support novel predictions. Participants rapidly learned novel transitions linking two previously separate sequences, allowing them to successfully anticipate upcoming environments on the new connected sequence immediately after learning. They continued to improve over the course of the experiment, suggesting both rapid and gradual updating of sequence representations. Multivoxel fMRI analyses revealed that hippocampus, but not visual cortex, exhibited increased pattern similarity for previously separate sequences that became connected vs. sequences that remained separate. Hippocampal integration emerged in early runs, suggesting rapid updating of sequence structure in the hippocampus. These integrated representations contained activity patterns from the initial sequences as well as new activity patterns not previously present, and predicted participants’ ability to update their predictions in behavior.

Our findings build on research demonstrating that memories for related experiences – typically, pairwise associations – become integrated in the hippocampus (Schlichting & Preston, 2015). These studies show that when one association (AB) shares features with another (AC), the indirectly related experiences (BC) become represented more similarly in the hippocampus, compared to unrelated pairs [12,19–21,25]. Such integration supports generalization and inferences about indirect relationships [46,47]. More broadly, memory integration may help form structured knowledge, creating internal models or “cognitive maps” that generalize across experiences to guide adaptive behavior [16–18], such as making predictions multiple steps into the future [3].

Unlike studies of pairwise associations that become linked, there are relatively few studies exploring hippocampal integration of large-scale, multistep associations. A recent study addressed this gap, demonstrating that the hippocampus constructs global environment representations that link separately learned routes to support flexible navigation [16]. Participants first learned to navigate within three separate routes in a virtual environment. After a one-day delay, they were taught that the routes were linked and then navigated across them to reach goal locations, requiring a global representation of the newly connected environment. Greater across-route hippocampal similarity *before* the environments were linked predicted more efficient navigation *after* they were linked [16]. Our findings extend this work by showing that integrated sequence representations in the hippocampus (1) emerge rapidly following new learning, (2) preserve stability with initial representations while incorporating new activity patterns, and (3) support flexible multistep predictions in newly connected sequences. Thus, hippocampal memory integration rapidly builds structured representations that guide flexible future-oriented behaviors.

We found that integrated hippocampal representations for the connected vs. unconnected sequence emerged in early runs, immediately after new learning. In contrast, there was no significant difference in pattern similarity between sequence conditions in late runs. We did not detect a significant *difference* between early and late runs, and therefore do not overinterpret the absence of integration in late runs; nevertheless, this pattern of results is somewhat puzzling. Behavioral performance on the Phase 2 Prediction Task continued to improve across runs, raising the question: why did across-map pattern similarity emerge immediately, but not increase further with practice? One possibility is that the integration signal we observed in early runs is partly due to effortful retrieval of the newly learned connections from the bridge environments. As these connections are used more often, it may become less effortful to remember and use them. For this explanation to account for our findings, however, this effortful application of the bridge “rule” must be implemented via a pattern change in the hippocampus that is consistent across the newly connected maps, because we did not observe any univariate activity differences in hippocampus between the connected vs. unconnected sequence conditions. A related possibility is that hippocampal integration is especially useful when task demands are high—such as during early runs, when participants must make multistep judgments across newly linked sequences. Bringing representations of the connected sequences closer together may allow the hippocampus to scaffold these difficult multistep judgments. With increased practice, however, participants may rely less on hippocampal integration to support performance, leading to improved behavior without corresponding increases in hippocampal pattern similarity. Complementary to this explanation, a recent study found that hippocampal differentiation of similar scene images occurred most prominently along a given feature dimension when that dimension most contributed to behavioral interference [48]. Together, this work and ours raise the possibility that hippocampal differentiation and integration mechanisms may be most pronounced when the task is more challenging for participants.

In addition to *when* integrated representations emerge, we also investigated the nature of these representations. We found that individual sequences maintained some stability with their initial representations even after they were updated into a newly connected sequence. However, there was no evidence for blending of the newly connected sequences. Instead, a partial correlation analysis revealed that across-map pattern similarity for the connected sequence was at least partly due to a novel pattern that was shared between the sequences in Phase 2, a pattern not present in either sequence originally, in Phase 1. This aligns with prior evidence that de novo representations support insight across related experiences [13]. In our experiment, most of each sequence remained unchanged from pre- to post-updating: the sequences were linked by two local connections, and the remaining connections were stable over time. Because the sequences retained some structure from the originally learned sequence, in addition to local changes at the bridge environments, it may be adaptive to retain partial similarity to the original neural representations while incorporating new information simultaneously. Supporting this idea, we found that across-map similarity was more pronounced in environments that could (vs. could not) reach the other map within 4 steps (the maximum distance for the correct “nearest environment” in our experiment) – indicating that representational change may be strongest in those parts of the sequence that have to undergo more updating. That said, we failed to find a significant *difference* in across-map similarity for environments that were within vs. not within 4 steps of the other map. Thus, this result requires additional investigation to be confirmed.

We observed evidence suggestive of integrated representations in the hippocampus for the connected sequences, but the nature of these memory traces is difficult to determine. One possibility is that the new, shared pattern that was not present in either sequence representation pre-integration (**Figure 5**) is indicative of the newly connected sequences becoming joined into a unified memory trace. An alternative possibility is that the memory traces for the sequences primarily remain separate, and the evidence for a novel, integrated pattern arises from a new memory link that accompanies co-activation of otherwise separate memory traces. It is not possible to tease apart these possibilities in the current design, and indeed this issue is a pervasive ambiguity in the literature on memory integration [49–53]. Determining which of these possibilities is underlying memory integration could be facilitated by studies using cellular-level imaging of memory traces pre- and post-integration.

Prior studies have shown important representational and functional differences between hippocampal subfields and between anterior and posterior hippocampus [14,54–57]. These differences extend to how these hippocampal regions represent overlapping events and sequences [12,56,58–64]. Our analyses were across the hippocampus as a whole because we did not have the spatial resolution to examine different hippocampal subfields. We also restricted our analyses to environment-selective voxels in hippocampus, further limiting our ability to examine subfield or long-axis differences. Future studies with high-resolution fMRI can test whether the integration we observed in our study was limited to certain regions of the hippocampus.

What kind of hippocampal learning mechanisms underlie sequence updating? Our prior work demonstrated that the hippocampus represents temporally extended sequences in a graded manner, with weaker representations for further-away environments [3]. These findings are consistent with computational accounts of successor representations, which propose that internal models cache predictions about successive states, with states weighted according to their distance [2,65,66]. A feature of successor representations, however, is that novel transitions are not easily incorporated, leading to predictions biased toward the original, pre-update sequence [66]. For example, after learning new transitions within a familiar sequence, predictive hippocampal activity continued to reflect the original successor state rather than the updated one [67]. Recent work suggests that successor representations can be modified more readily in some circumstances, particularly via replay – with even minimal on-task replay sufficient to incorporate novel transitions [68]. In our study, replay between runs may have allowed participants to adjust cached representations, integrating the newly learned transitions to support accurate multistep predictions.

An alternative account is that participants engaged a model-based strategy, explicitly representing each successive link between environments [69]. Unlike successor representations, model-based learning readily supports rapid updating because it explicitly stores information about state-to-state transitions. When new transitions are encountered, the transition model can be directly revised, allowing predictions to be recomputed online from the updated structure [66]. We have found evidence consistent with such a link-based strategy that supports multistep predictions particularly after memory consolidation [36]. Hippocampal representations may flexibly shift between cached successor representations and model-based computations depending on task demands. Future work could disentangle these mechanisms by testing how replay, consolidation, and task demands interact to support the balance between successor representations and model-based learning in hippocampal sequence representations.

In summary, we show that new learning rapidly integrates sequence representations in the hippocampus. The extent to which sequences were integrated was related to the behavioral ability to update multistep predictions. Although participants quickly learned the new sequence structure, their ability to make predictions about upcoming environments across sequences also improved with time, implying both rapid and gradual updating of learned representations. This opens the door for future work investigating how internal models are both rapidly and gradually updated in the brain. Thus, our work sheds light on how and when internal models are represented and updated, and how they are used to guide flexible and adaptive behavior.

## CRediT Statement

**Hannah Tarder-Stoll**: Conceptualization, Data Curation, Investigation, Methodology, Project Administration, Software, Writing—original draft, Writing—review and editing; **Chris Baldassano**: Conceptualization, Methodology, Funding Acquisition, Project Administration, Resources, Supervision, Writing—original draft, Writing—review and editing; **Mariam Aly**: Conceptualization, Methodology, Funding Acquisition, Project Administration, Resources, Supervision, Writing—original draft, Writing—review and editing.

## Acknowledgements

This work was funded by a National Institutes of Health Research Project Grant (R01EY034436) and a Zuckerman Institute Seed Grant for MR Studies (CU-ZI-MR-S-0016) to M.A. and C.B. We would like to thank the Alyssano Group for helpful advice on this project.

