## Supplementary Materials for "The Hippocampus Rapidly Integrates Sequence Representations During Novel Multistep Predictions"

An example of a video of the bridge environment in Map A transitioning to Map B, and from Map B back to Map A, on the connected sequence can be here. In this example, the connected sequence was the green path.
